# Computational saturation mutagenesis of SARS-CoV-1 spike glycoprotein: stability, binding affinity, and comparison with SARS-CoV-2

**DOI:** 10.1101/2021.06.30.450547

**Authors:** Adebiyi Sobitan, Vidhyanand Mahase, Raina Rhoades, Dejaun Williams, Dongxiao Liu, Yixin Xie, Lin Li, Qiyi Tang, Shaolei Teng

## Abstract

Severe Acute respiratory syndrome coronavirus (SARS-CoV-1) attaches to the host cell surface to initiate the interaction between the receptor-binding domain (RBD) of its spike glycoprotein (S) and the human Angiotensin-converting enzyme (hACE2) receptor. SARS-CoV-1 mutates frequently because of its RNA genome, which challenges the antiviral development. Here, we performed computational saturation mutagenesis of the S protein of SARS-CoV-1 to identify the residues crucial for its functions. We used the structure-based energy calculations to analyze the effects of the missense mutations on the SARS-CoV-1 S stability and the binding affinity with hACE2. The sequence and structure alignment showed similarities between the S proteins of SARS-CoV-1 and SARS-CoV-2. Interestingly, we found that target mutations of S protein amino acids generate similar effects on their stabilities between SARS-CoV-1 and SARS-CoV-2. For example, G839W of SARS-CoV-1 corresponds to G857W of SARS-CoV-2, which decrease the stability of their S glycoproteins. The viral mutation analysis of the two different SARS-CoV-1 isolates showed that mutations, T487S and L472P, weakened the S-hACE2 binding of the 2003-2004 SARS-CoV-1 isolate. In addition, the mutations of L472P and F360S destabilized the 2003-2004 viral isolate. We further predicted that many mutations on N-linked glycosylation sites would increase the stability of the S glycoprotein. Our results can be of therapeutic importance in the design of antivirals or vaccines against SARS-CoV-1 and SARS-CoV-2.

**Author Summary:** Severe acute respiratory syndrome coronavirus (SARS-CoV-1) is an RNA virus that undergoes frequent mutations, which may result in more virulent SARS-CoV-1 variants. To prevent another pandemic in the future, scientists must understand the mechanisms of viral mutations and predict if any variants could become a dominant. The infection of SARS-CoV-1 in cells is largely depending on the interactions of the viral Spike (S) and human angiotensin-converting enzyme 2 (hACE2). We applied a computational method to predict S missense mutations that will make SARS-CoV-1 more virulent. We are interested in the variants that can change SARS-CoV-1 spike protein stability and/or change the virus-receptor interactions. We mutated each residue of SARS-CoV-1 spike to all possible amino acids; we calculated the differences between the folding energy and binding energy of each variant and the wildtype and identified the target S mutations with significant effects on protein stability and protein-protein interaction. We found some viral mutations could destabilize S and weaken S-hACE2 binding of SARS-CoV-1 isolate. Our results show that the computational saturation mutagenesis is a reliable approach in the analysis and prediction of missense mutations.

## Introduction

The severe acute respiratory syndrome coronavirus (SARS-CoV-1) belongs to a family of coronaviridae that are enveloped, positive-strand RNA viruses (1). In November 2002, the first case of SARS-CoV-1 occurred in the Guangdong province in China. The symptoms include upper respiratory infections, fever, chills, and general body weakness (2). The other signs showing human-to-human transmission were coughing and sneezing. Horseshoe bat species might be the origin due to their sequence similarity (3). By the end of the SARS epidemic in 2003, SARS had spread to over two dozen countries resulting in more than 8000 laboratory-confirmed cases and approximately 800 deaths (4). The recent outbreak of a newer strain of coronavirus, SARS-CoV-2, began in December 2019 in the Wuhan city in China (5). In a month, this new coronavirus had spread across the world due to global travels. Compared to SARS-CoV-1, SARS-CoV-2 has a higher infection rate. As of June 6, 2021, the number of confirmed global cases and global deaths due to SARS-CoV-2 are ~173 million and ~3.7 million, respectively (6).

The Spike protein (S) is a structural protein that protrudes outwards from the virus surface. The role of S is to mediate viral entry into the host’s cells. Structural studies of SARS-CoV-1’s S revealed the presence of two subunits, S1 (residues 12-667) is in the N-terminal and S2 (residues 667-1190) is in the C-terminal. Studies of mammalian coronaviruses with similarity to the SARS-CoV-1 showed that the S1 subunit helps with hACE2 receptor attachment, while the S2 subunit helps with the fusion (1). Expression analysis showed that a fragment of the S1 subunit, the receptor-binding domain (RBD), residues 306 - 527, is enough for tight binding to the human Angiotensin-converting enzyme 2 (hACE2) receptor (7). A shorter fragment, residues 424 - 494, within the RBD interacts with the hACE2 receptor in humans. This fragment, the receptor-binding motif (RBM), forms a loop that fits perfectly into the peptidase domain (PD) of the hACE2 receptor. The SARS-CoV-1 glycoprotein has two cleavage sites that promote viral infection. The first cleavage site is in the 667-668 residue positions. As the virus enters the host’s cell, its spike protein is cleaved into the S1 and S2 subunit by protease activity. The second cleavage site is in the 797-798 residue positions. Cleavage at this position detaches the fusion protein from the S2 subunit and allows the fusion protein to bind with the host’s membrane (8).

One similarity shared by SARS-CoV-1 and SARS-CoV-2 is that they use hACE2 as the receptor to enter human cells [10, 12]. Both SARS-CoV-1 and SARS-CoV-2 S proteins bind to the Peptidase domain in the N-terminal of the hACE2 receptor (1). The other domain, the Collectrin domain, is found in the C-terminal of the hACE2 receptor. However, studies showed they exhibit varying binding affinities. A recent study reported that SARS-CoV-2 S protein has a 10-to-20 fold higher affinity to hACE2 than that of SARS-CoV-1 (7). The interaction between the S protein of SARS-CoV-1 and the hACE2 receptor initiates entry into the human cell [12]. The higher affinity SARS-CoV-2 has for hACE2 may explain the virulent nature of its infection (10). Another similarity is the sequence and structural homology between the S proteins of SARS-CoV-1 and SARS-CoV-2. However, despite the similarities, a study evaluated the binding of SARS-CoV-2 to experimentally verified monoclonal antibodies (mAbs) against SARS-CoV-1. The result showed a slight contrast in cross-reactivity, which had no binding between SARS-CoV-2 and the three mAbs (11). This result supports the hypothesis that the slight difference in their sequences/structures might be responsible for the varying infectivity between SARS-CoV-1 and SARS-CoV-2. In a previous study, we indicated that SARS-CoV-2 has a stronger affinity towards hACE2 than SARS-CoV-1 because of its higher electric field density (12). hACE2 plays a role in the renin-angiotensin pathway, that it maintains cardiovascular homeostasis (13). hACE2 participates in microbial infection by serving as an entry point for coronaviruses (14). The variation of hACE2 across species explains why SARS-CoV-1 infects humans and not rats nor mice. A study manipulated the protein sequences of the hACE2 of rats and mice by mutating specific residues to the residues in humans. The result found an increase in infectivity when the mouse or rat hACE2 has human residues in certain positions (15). Understanding the interactions between the contact residues of SARS-CoV-1 and hACE2 can provide insights into how SARS-CoV-2 enters human cells.

A recent bioinformatic study on SARS-CoV-1 revealed that different subdomains of the spike protein exhibit varying mutation rates. These varying mutation rates might be the driving force for the difference in the infectivity rate of SARS-CoV-1 (16). Computational analysis has proven an effective method in studying protein dynamics (17). SARS-CoV-1 has an unstable RNA genome. Comparatively, experimental approaches are limited in their ability to generate functional measurements of all potential missense mutations, which would be expensive, laborious, and time-consuming. Recently, we used a computational approach to predict and analyze missense mutations in SARS-CoV-2 S protein (18). We predicted several missense mutations that affect the stability and binding affinity of SARS-CoV-2. These target mutations include D614G, N501Y, and K417N found in South Africa, United Kingdom, and Brazil variants. As a result, we employed the bioinformatics tools to analyze the effects of missense mutations on the stability and protein-protein interactions of SARS-CoV-1. This approach is fast and less tedious in identifying key residues, which will help design therapeutic drugs against SARS-CoV-1.

## Results

### Sequence and structural alignments of S proteins of SARS-CoV-1 and SARS-CoV-2

The Jalview tool (19) shows the aligned residues, the quality of the alignment, the conservations scores, and the consensus between the RBD sequences of SARS-CoV-1 and SARS-CoV-2 (Fig 1A). The sequence alignment using clustal omega algorithm within the Jalview tool indicates a ~76% sequence identity between RBD regions of SARS-CoV-1 and SARS-CoV-2. The bright yellow bars represent high conservation and quality between the aligned residues within each column. As shown in Fig 1B, the structural alignment revealed the evolutionary relationship between SARS-CoV-1 and SARS-CoV-2. SARS-CoV-1 S (PDB ID: 6ACG, chain A) aligned with SARS-CoV-2 S (PDB ID: 6VYB, chain A) with an RMSD of 2.272. Furthermore, we performed the structural alignment of the RBDs of SARS-CoV-1(PDB ID: 2AJF, chain E) and SARS-CoV-2 (PDB ID: 6M17, chain E), and it yielded an RMSD of 1.043. We selected a shorter fragment, RBM, from the RBD and performed the structural alignment of the RBM of both coronaviruses. This alignment yielded an RMSD of 0.878, which is more homologous. The shape and spatial orientation of the structural alignments overlapped, which indicates close atomic coordinates between the two structures.

**Fig 1:**
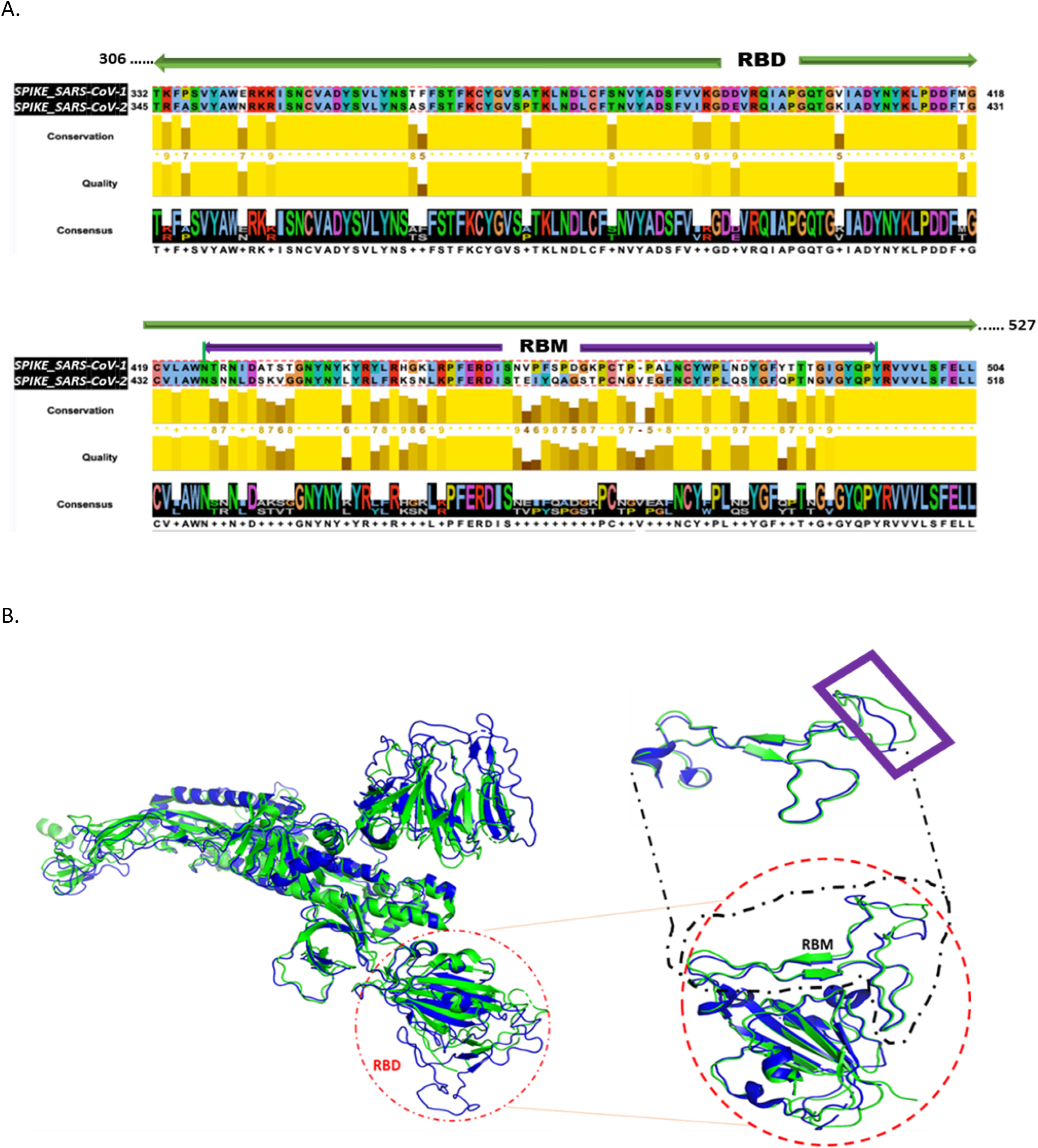
Sequence and Structural Alignment of SARS-CoV-1 S and SARS-CoV-2 S. (A) Sequence alignment of the receptor binding domain (RBD) sequences (B) Structural alignment. *Left*: Alignment of the full-length spike proteins of SARS-CoV-1 (Blue) and SARS-CoV-2 (Green). Red circle indicates orientation of RBD. *Right*: Alignment of RBDs. Black dashed square represents the alignment of the RBMs. Purple rectangle reveals regions with imperfect alignment.

### Effects of mutations on full-length SARS-CoV-1 S stability (ΔΔG)

The SARS-CoV-1 S protein has 1255 residues that were used to generate 23,845 non-redundant missense mutations. The effect of each mutation on the stability of the SARS-CoV-1 S protein was evaluated. Of the total mutations performed, 20,083 missense mutations generated energy changes, while the remaining 3,762 missense mutations gave no output due to missing residues on the protein structure.

Fig 2A shows that 11,635 of the 20,083 (58%) missense mutations increased the free energy of the S protein by at least 0.5 kcal/mol, 2,964 of 20,083 (15%) missense mutations reduced the S protein’s free energy by at most −0.5 kcal/mol, and 5,484 of 20,083 (27%) had a neutral effect on the stability of the wildtype S protein. The standard error of energies calculation using the FoldX suite is ~0.5kcal/mol (20). Therefore, the folding energy changes within the range (−0.5<ΔΔG <0.5) are insignificant or categorized as having neutral effect. In more specific categories, 4,767 mutations had a highly destabilizing effect (ΔΔG>2.5kcal/mol) on the spike protein, 6,868 mutations moderately destabilize the spike’s protein (0.5<ΔΔG≤2.5kcal/mol), 5484 mutations had a neutral effect (−0.5<ΔΔG≤0.5kcal/mol), 2,816 mutations moderately stabilize the S protein (−2.5ΔΔG≤−0.5kcal/mol), and 148 mutations have a highly destabilizing effect (ΔΔG<−2.5kcal/mol) on the S protein.

**Fig 2.**
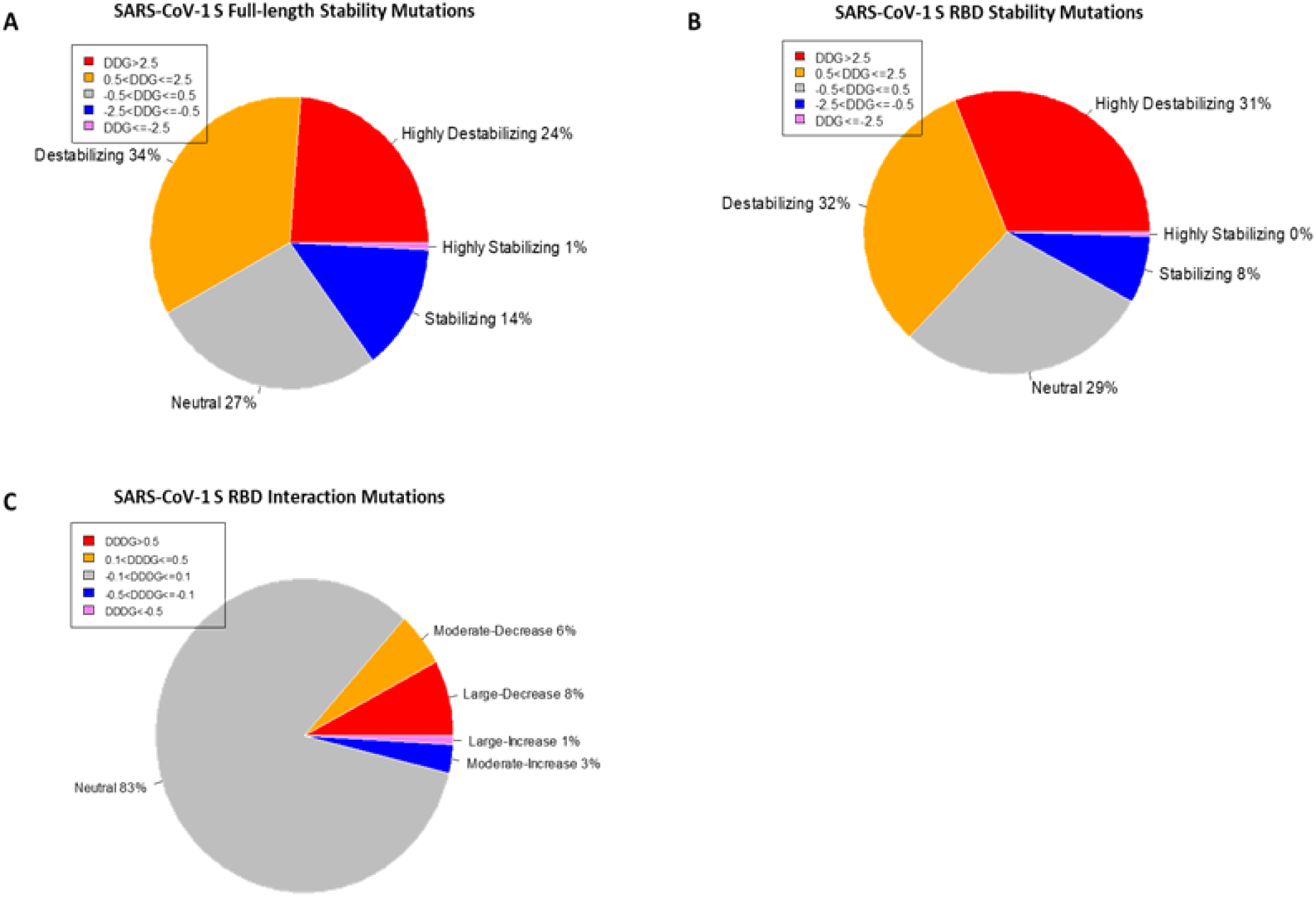
Distribution of the effects of missense mutations on protein stability and binding affinity. Pie charts of the effects of missense mutations on SARS-CoV-1 S (full-length) stability (A) and S RBD stability (B). (C) Pie chart of the effects of missense mutations on SARS-CoV-1 S RBD binding affinity.

The line chart and the overall heatmap show the distribution of the mutations along the entire length of the spike protein (Fig 3A). The red lines represent the positive mean values while the blue lines represent the negative mean values. The bubbles represent the folding energy changes when all the residues were mutated to Alanine. Based on the average of ΔΔG, mutations with the highest destabilizing effects were found in positions 839, 634, 418, 430, 536, and 1113 (Fig 3). The two most destabilizing missense mutations, A430W and A430F, cause energy changes at 66.18kcal/mol and 56.4kcal/mol, respectively. This position is also within the RBM of the SARS-CoV-1 spike S protein. On the other hand, mutations with the highest stabilizing effects were found in positions: 1059, 981, 500, 1089, 150, and 247 (Fig 3). The two most stabilizing missense mutations, G981W and T1059F, reduced the free energy of the wildtype structure by −5.16 kcal/mol and −4.98 kcal/mol, respectively. The residue positions with the highest and lowest mean folding energy changes (ΔΔG) were G839 and T1059, which are both within the S2 subunit of the S protein. Compared to the mean values of all mutations, the missense mutations to Alanine had more destabilizing effects than stabilizing effects. The two “white” gaps in the overall heatmap represent missing values in the residue positions 661 – 673 and 812 – 831, respectively.

**Fig 3:**
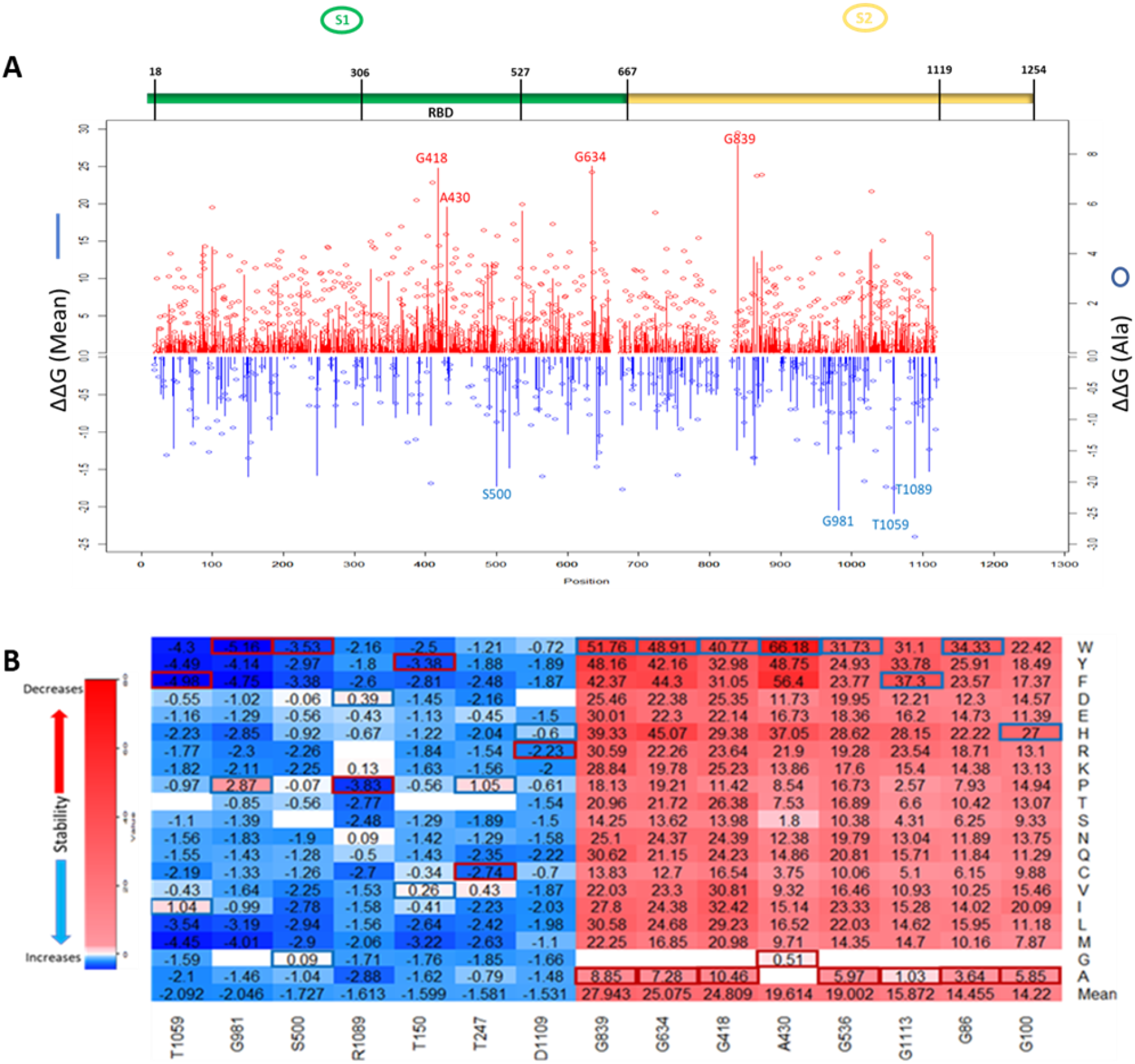
Visualization of ΔΔG values caused by missense mutations on SARS-CoV-1 S (full-length). (A) Line chart showing mean ΔΔG values of destabilizing mutations (red upward lines) and stabilizing mutations (Blue downward lines). Red and Blue bubbles show ΔΔG values of Alanine as alternate residues (Top). Overall heatmap of all missense mutations showing different regions/domains (Bottom). (B) Heatmap of top 5 destabilizing mutations and top 5 stabilizing mutations on SARS-CoV-1 S (full-length). Blue rectangle represents maximum ΔΔG values. Red rectangle represents minimum ΔΔG values.

### Effects of mutations on SARS-CoV-1 S RBD stability (ΔΔG)

We utilized the crystal structure of the RBD of SARS-CoV-1 (PDB ID:2AJF) for computational prediction of the effect of computed mutations on the stability of SARS-CoV-1 RBD. This analysis gave an alternative perspective on the predictive power of computational tools. We generated 3,841 mutations and calculated the folding energy change (ΔΔG) caused by each missense mutation. Like the full spike analysis, 63% of the missense mutations destabilized the SARS-CoV-1 RBD structure. Meanwhile, 8% of the missense mutations stabilized the SARS-CoV-1 RBD structure. Fig 2B shows the pie chart of the effects of the missense mutations on stability of SARS-CoV-1 S RBD. The distribution of the effect of all missense mutations in both the RBD and the full-length S stability analysis correlates (r = 0.6824). Observation of individual mutations revealed analytic similarity in both stability analysis. The heatmaps in Fig 3B and 4A also show similarity in missense mutations with large effects on the stability of the SARS-CoV-1 S stability. For example, the missense mutation A430W had a large effect on both the stability of the RBD (Fig 4B) and the entire spike protein of the SARS-CoV-1.

**Fig 4.**
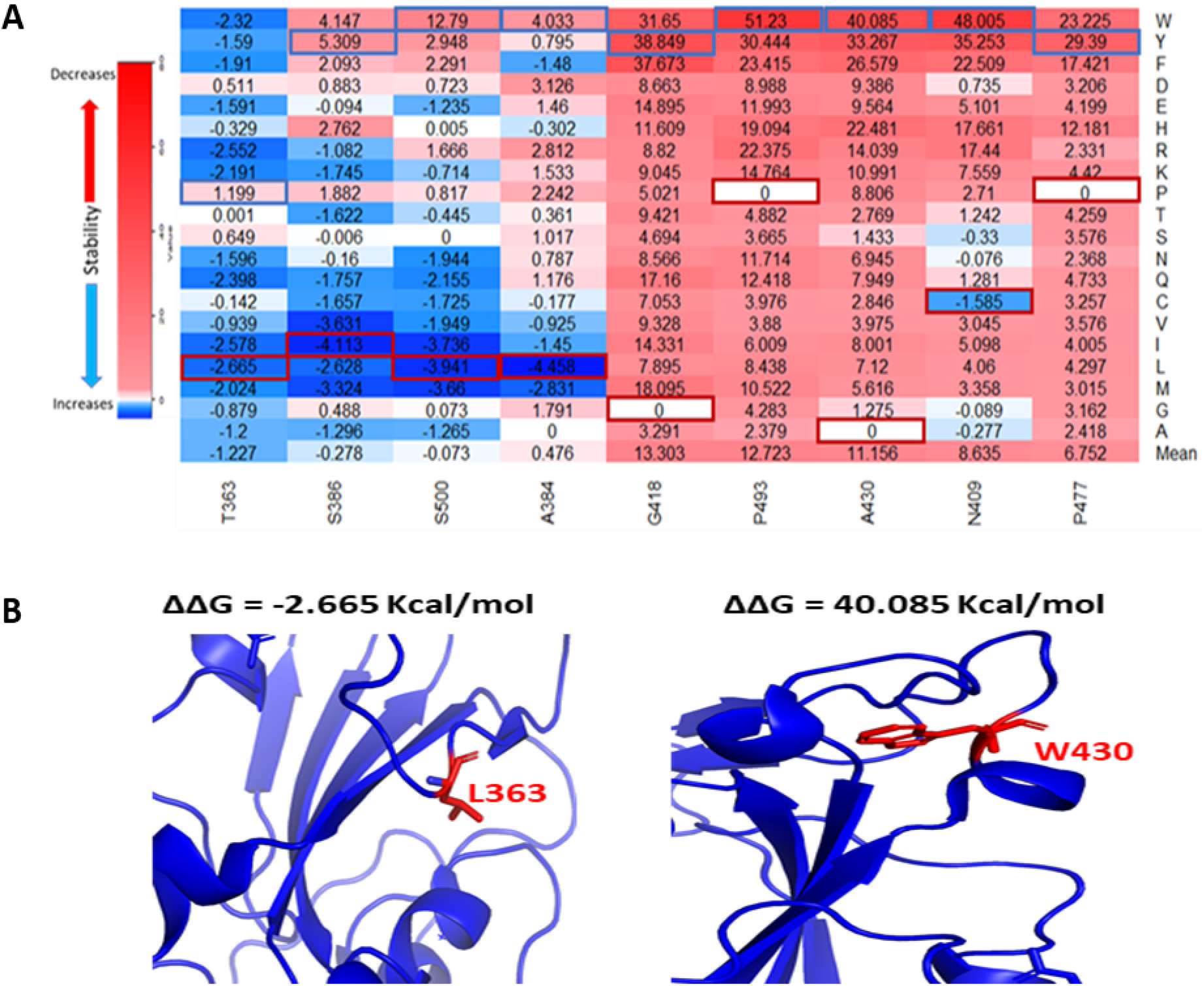
Effects of the target missense mutations on SARS-CoV-1 S RBD protein stability. (A) Heatmap of top destabilizing mutations and top stabilizing mutations (ΔΔG) Values up the scale(red) decreases stability/binding affinity and vice versa (blue). The blue rectangular box represents maximum values. The red rectangular box represents minimum values. (B) Structural representation of SARS-CoV-1 S RBD showing stabilizing (T363L) and destabilizing (A430W) mutations in red and their side chains.

### Comparison of effects of mutations on stabilities of full-length S proteins of SARS-CoV-1 and SARS-CoV-2

We compared the mutational effects of seven residues on SARS-CoV-1 S protein to corresponding residues on SARS-CoV-2 S protein. Four residues (T1059, G981, S500, and R1089) had the highest mean stabilizing effects, while three residues (G839, G634, and A430) had the highest mean destabilizing effects. Pairwise alignment of SARS-CoV-1 S and SARS-CoV-2 S showed that the residues T1059, G981, S500, R1089, G839, G634, and A430 on SARS-CoV-1 corresponds with residues T1077, G999, S515, R1107, G857, G648, and S443 on SARS-CoV-2, respectively. As shown in Table 1, the missense mutations on SARS-CoV-1 S protein had similar effects on its stability when compared to corresponding residues on SARS-CoV-2 S protein. We observed that the missense mutation A430S destabilizes SARS-CoV-1 S protein by introducing the folding energy change at 1.8 kcal/mol. Consistently, S443A in corresponding position of SARS-CoV-2 increase the S stability (ΔΔG = −0.768 kcal/mol). Interestingly, SARS-CoV-1 D600G can destabilize S (ΔΔG= 0.21 kcal/mol). D614G, the dominant variant of SARS-CoV-2, corresponds to D600G but was predicted to stabilize S (ΔΔG= −0.784 kcal/mol). In position 247, the substitution of Threonine with Alanine increased the stability of the SARS-CoV-1. Because the corresponding residue on SARS-CoV-2 was missing, we extrapolated that the presence of Alanine on position 260 on the SARS-CoV-2 would increase its stability.

**Table 1:** Mapping and comparing the effects of SARS-CoV-1 and SARS-CoV-2 missense mutations on protein stability (**Top**) and binding affinity (**Bottom**).

| SARS-CoV-1 |  |  | SARS-CoV-2 |  |  |
| --- | --- | --- | --- | --- | --- |
| Protein Stability |  |  |  |  |  |
| MUTATION | $\Delta\Delta G$<br>(kcal/mol) | Effect | MUTATION | $\Delta\Delta G$<br>(kcal/mol) | Effect |
| G839W | 51.76 | Decrease | G857W | 58.212 | Decrease |
| G839Y | 48.16 | Decrease | G857Y | 43.523 | Decrease |
| G634W | 48.91 | Decrease | G648W | 43.326 | Decrease |
| A430W | 66.18 | Decrease | S443W | 33.41 | Decrease |
| A430F | 56.4 | Decrease | S443F | 22.173 | Decrease |
| A430S | 1.8 | Decrease | S443A | -0.768 | Increase |
| A430Y | 48.75 | Decrease | S443Y | 25.052 | Decrease |
| D600G | 0.21 | Decrease | D614G | -0.784 | Increase |
| T1059F | -4.98 | Increase | T1077F | 3.263 | Decrease |
| G981W | -5.16 | Increase | G999W | 27.099 | Decrease |
| G981F | -4.75 | Increase | G999F | 25.374 | Decrease |
| S500W | -3.53 | Increase | S514W | -3.303 | Increase |
| T247C | -2.74 | Increase | A260C | NA | NA |
| T247A | -0.79 | Increase | A260T | NA | NA |
| Binding Affinity |  |  |  |  |  |
| MUTATION | $\Delta\Delta\Delta G$<br>(kcal/mol) | Effect | MUTATION | $\Delta\Delta\Delta G$<br>(kcal/mol) | Effect |
| G488P | 15.921 | Decrease | G502P | 11.767 | Decrease |
| T487Y | 20.018 | Decrease | N501Y | 4.55 | Decrease |
| T487N | 2.284 | Decrease | N501T | -1.346 | Increase |
| T487W | 13.42 | Decrease | N501W | 3.118 | Decrease |
| G482E | 5.758 | Decrease | G496E | 7.278 | Decrease |
| S432Y | -1.686 | Increase | V445Y | -0.253 | Increase |
| S432V | -0.22 | Increase | V445S | -0.0045 | Neutral |
| N479M | -1.42 | Increase | Q493M | -0.3 | Increase |
| N479Q | -0.607 | Increase | Q493N | 0.679 | Decrease |
| P462D | -0.97 | Increase | A475D | 0.633 | Decrease |
| P462A | 0.391 | Decrease | A475P | 0.282 | Decrease |

### Effects of mutations on SARS-CoV-1 S RBD binding affinity (ΔΔΔG)

The protein structure (PDB ID:2AJF) used for interaction analysis covered the S RBD that interact with the hACE2 receptor in humans. This RBD chain contains 180 residues, which spanned from residual position 323 to 502. A total of 3,420 mutations were computed, with 114 missing values. The remaining 3,306 mutations were classified into one of five categories according to their binding energy changes (ΔΔΔG). As shown in Fig 2C, 257 mutations had a large-decrease effect, 183 mutations had a moderate-decrease effect, 2,738 mutations had a neutral effect, 97 mutations had a moderate-increase effect, and 31 mutations had a large-increase effect. Fig 6E shows the mutations with the largest effects, based on ΔΔΔG mean values from all possible mutations in that position.

The seven topmost mutations with decreasing effects are found in positions which have mean values greater than 1.5 kcal/mol. While the six topmost mutations with increasing effects are found in positions which have mean values less than −0.3 kcal/mol. The values in red boxes represent the minimum values in each position, while the values in blue boxes represent maximum values. The overall distribution of the missense mutations as they change ACE2 binding affinity of the SARS-CoV-1 S protein can be seen in Fig 5. There are 7 distinct regions or domains in the heatmap: [337-338], [388-396], [402-409], [424-436], [439-447], [460-467], and [472-494]. Four of the seven distinct regions are concentrated within the RBM, and they span longer stretches of residues. Most of the red and blue spikes on the line chart are also within the RBM. These distinct regions can provide insight into which residues play a key role in the interaction between SARS-CoV-1 S protein and the hACE2 receptors in humans.

**Fig 5.**
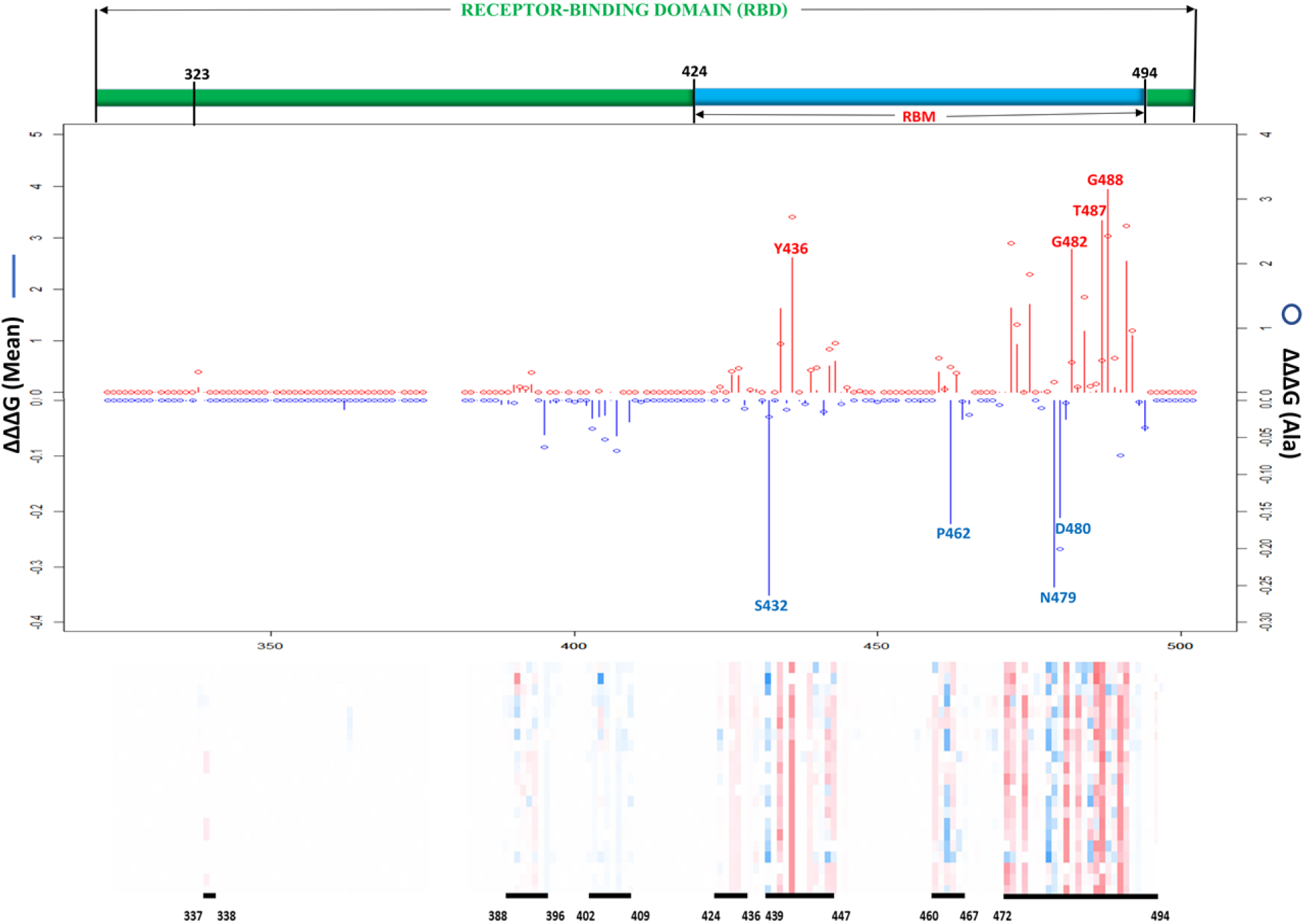
Effects of all possible missense mutations on SARS-CoV-1 S RBD binding affinity. Line chart showing mean ΔΔΔG values of mutations decreasing binding affinity (red upward lines) and mutations increasing binding affinity (Blue downward lines). Red and Blue bubbles show ΔΔG values of Alanine as alternate residues (Top). Overall heatmaps showing seven hot regions/domains and residue span (Below).

### Analysis of interaction between SARS-CoV-1 and hACE2 receptor

The contact residues between hACE2 and the SARS-CoV-1 S were shown in Fig 6A. Not all the residues within the PD interact with SARS-CoV-1 and vice-versa. However, mutations on interacting or contact residues between hACE2 and SARS-CoV-1 showed to significantly affect their binding affinities. Fig 6 B-D shows key residues at the interface of hACE2 and SARS-CoV-1 that contributes to the binding affinity between the two molecules. The effect of mutation on the binding energy of T487 of the S protein weaken its affinity for residues, Y41 and K353, on hACE2 and vice-versa.

**Fig 6.**
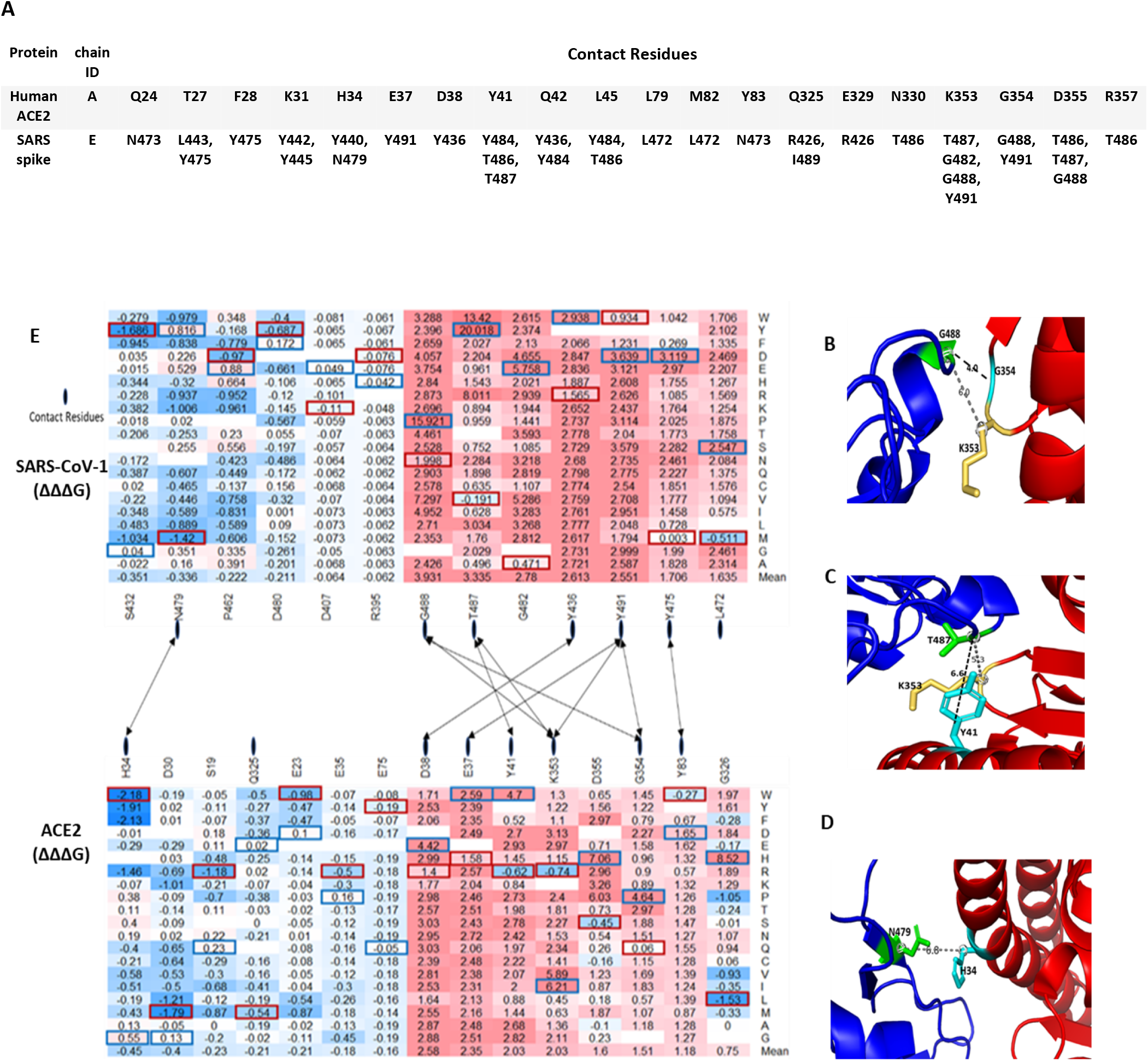
Top missense mutations on SARS-CoV-1 S RBD-ACE2 interface. (A) Contact residues between Human ACE2 (Top) and SARS-CoV-1 S RBD protein (Bottom). (B, C, D) Structural representation of important binding residues (SARS-CoV-1 S RBD interacting residue in green and hACE2 interacting residues in yellow and cyan). (B) RBD G488 interacts with K353 and G354 on ACE2. (C) RBD T487 interacts with Y41 and K353 on ACE2. (D) RBD N479 interacts with H34 on ACE2. Dotted lines indicate distance between two residues. (E) Heatmap of the ΔΔΔG of target S RBD mutations. Dotted lines indicate contact residues between the SARS-CoV-1 S RBD and hACE2.

### Comparison of effects of mutations on the binding affinities of SARS-CoV-1 and SARS-CoV-2 S protein

As shown in Table 1, Asparagine is found on position 501 in SARS-CoV-2 which corresponds to Threonine on position 487 in SARS-CoV-1. The computed missense mutation T487N increased the ΔΔΔG by 2.28 kcal/mol and decreased the ΔΔG by −0.14. In contrast, N501T, on the SARS-CoV-2 decreased ΔΔΔG by −1.346kcal/mol. Tyrosine and Tryptophan have nonpolar aromatic chains therefore the missense mutations T487Y and T487W, had similar effects (decrease binding affinity) on corresponding residue on SARS-CoV-2, N501Y and N501W, respectively. Most of the missense mutations on positions S432, P462, and N479, on the average, increase the binding affinity of SARS-CoV-1. However, the corresponding residues on SARS-CoV-2 are Valine, Alanine, and Glutamine, respectively. The effect of missense mutations on these residues were evaluated by examining missense mutations on corresponding positions on SARS-CoV-1. However, missense mutation N479Q decreased the ΔΔΔG by −0.607 kcal/mol and increased ΔΔG by 1.91 kcal/mol. A possible explanation is that the missense mutations created Van der Waal clashes with neighboring atoms because of shorter distance; The missense mutation, T487Y, caused a differential Van der Waals of −0.449kcal/mol, which led to a greater repulsion from nearby atoms. Recent SARS-CoV-2 variant L452R (ΔΔΔG= −0.395kcal/mol; ΔΔG = 0.021kcal/mol) corresponds to SARS-CoV-1 K439R (ΔΔΔG= 0.247kcal/mol; ΔΔG= 0.41 kcal/mol). The change in residue from Lysine in SARS-CoV-1 S protein to Leucine in SARS-CoV-2 S protein may be responsible for the increase in binding affinity caused by L452R.

### Effects of mutations on Post-Translational Modification sites of S protein

Post-Translational modifications (PTMs) of the SARS-CoV-1 S protein are responsible for the folding, maturation, and function of the S protein. An important PTM utilized by SARS-CoV-1 is the O- and N-linked Glycosylation, which plays a key role in the shielding of viruses from the host’s immune system (21). As a result, viruses evolve to become glycosylated as much as possible (22). One O-linked glycosylation site, S336, located within the RBD was predicted using online webserver (23). On the average, the computed missense mutations in this position increase the stability of SARS-CoV-1. For example, the missense mutation, S336E, reduces the ΔΔG by −1.93 kcal/mol, thereby stabilizing the SARS-CoV-1 spike protein. Conversely, S336V increases the ΔΔG by 1.28 kcal/mol, thereby destabilizing the SARS-CoV-1 structure. We used five online prediction tools in our N-linked glycosylation sites prediction-NetNGly 1.0 (24), N-GlyDE (25), SPRINT-Gly (26), Glycopp v1.0 (27), and Glycopred (28). We narrowed our results to sites predicted by at least two or three prediction tools. Of these, we selected 20 N-linked glycosylation sites located within the spike protein of the SARS-CoV-1. Three of these, N318, N330, N337, were located within the RBD of SARS-CoV-1 spike protein. On the average, mutations at position N318 were predicted to increase the stability of SARS-CoV-1. However, on the average, the mutations on N330 and N337 were predicted to have neutral effects.

Fig 7 shows the target mutations in twenty N-linked glycosylation sites, one O-linked glycosylation site, and one palmitoylation site. Palmitoylation modifies the spike protein through cysteine-rich residues (29). This is known to mediate the fusion of the spike protein to hACE2. Predicted palmitoylation sites (30) were found in positions C19, C1217, C1218, C1222, C1232, C1235, and C1236. The average computed mutations at the C19 position predicted neutral effects, which showed no significant effect in the stability of SARS-CoV-1. However, the remaining six palmitoylated were located outside the residues covered by the structural protein.

**Fig 7.**
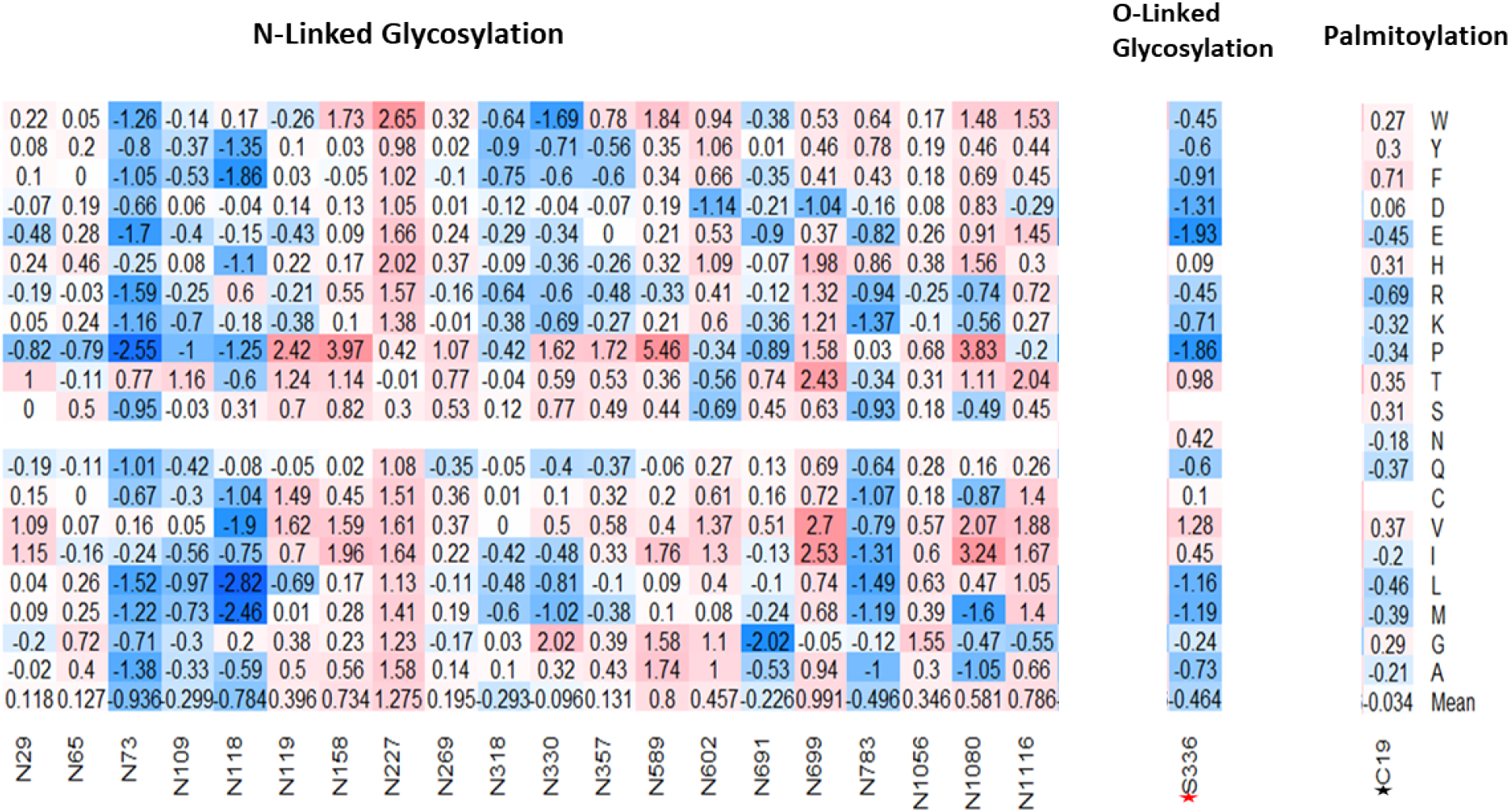
Effects of missense mutations on putative post translational sites on SARS-CoV-1 full-length S protein. (Left-Right) 20 N-Glycosylation sites, 1 O-Glycosylation site (S336), and 1 Palmitoylation site (C19).

### Statistical analysis of SARS-CoV-1 S protein mutation pathogenicity

We used sequence-based mutation pathogenicity tools to predict the damaging effect of our computed mutations on the SARS-CoV-1 S function. We analyzed 25,101 missense mutations using the full-length SARS-CoV-1 S (1 – 1255). The PolyPhen2 scores gave probabilistic values on the tolerance and deleterious effect of missense mutations. A score less than 0.446 is considered benign, a score greater than 0.446 but less than 0.908 is considered possibly damaging, and a score greater than 0.908 is considered probably damaging. In Fig 8A, the missense mutations with neutral effect were predicted to be mostly tolerated with some classified as benign, while the mean value, as shown by the red line, is considered possibly damaging. Whereas the mean values of the moderately increasing and decreasing mutations were predicted to be possibly damaging. However, the mean values of large increasing and decreasing mutations were predicted to be probably damaging. The analysis of variance (ANOVA) showed that the means of all five categories were significantly different, with P-value < 2e-16 (Fig 8A).

**Fig 8.**
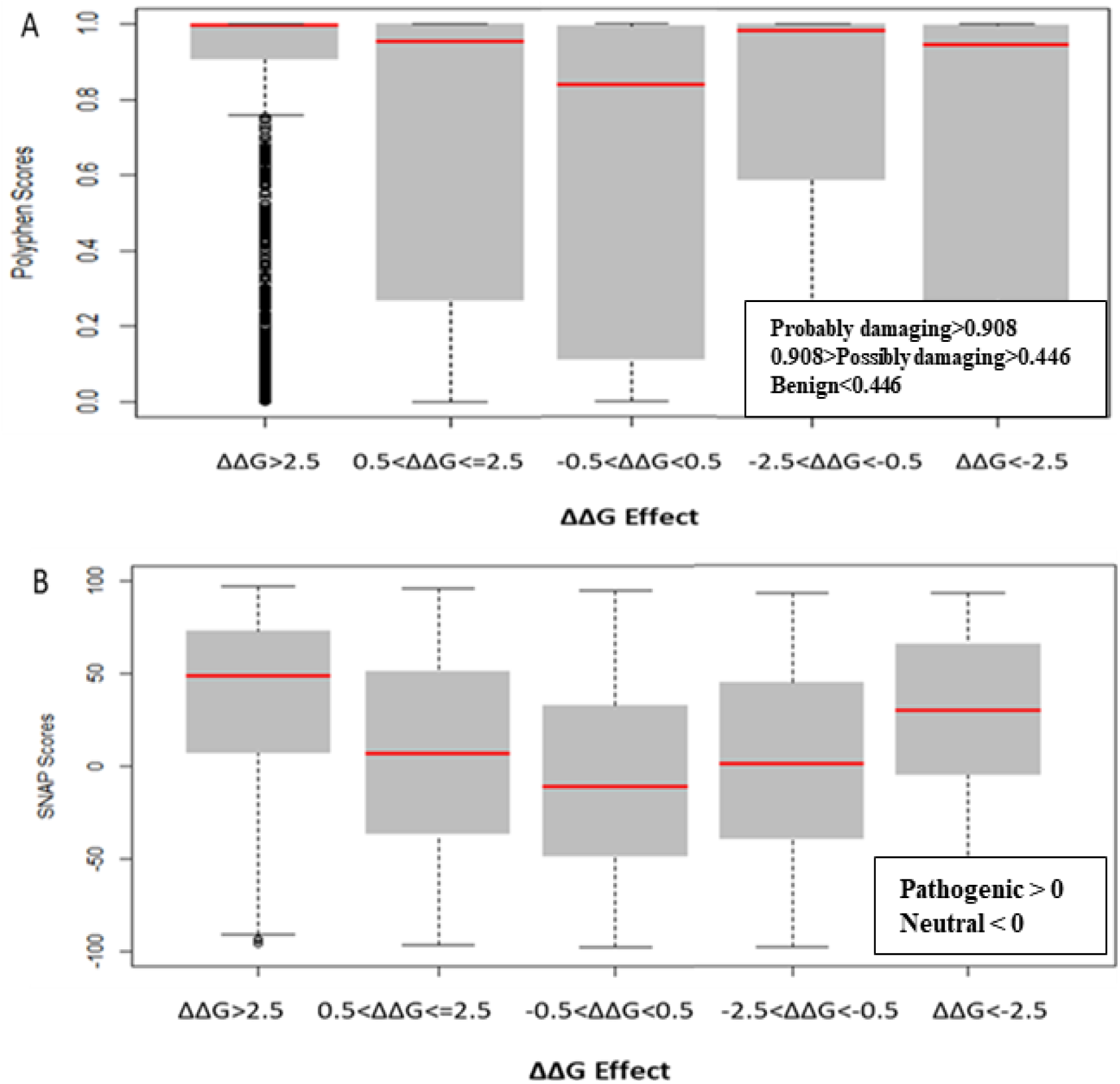
Boxplots of the prediction of mutation pathogenicity on full-length Spike. (A) PolyPhen2 scores and (B) SNAP scores against five categories of the effects of mutations on SARS-CoV-1 full-length S protein stability.

SNAP is a neural network machine learning algorithm that accepts protein structure as input for functional predictions (31). SNAP scores less than zero have neutral effect, while SNAP scores greater than zero have a pathogenic effect. 13,179 (~53%) of the total 25,101missense mutations have pathogenic effects. We performed further analysis to correlate the effects of folding energy change ΔΔG on stability with their SNAP scores. As shown in Fig 8B, the SNAP scores of missense mutations with effects greater than 2.5 kcal/mol or less than −2.5 kcal/mol were higher than missense mutations with moderate effects (0.5<ΔΔG <= 2.5 or −2.5 =< ΔΔG < −0.5 kcal/mol). Furthermore, the missense mutations with neutral effects on SARS-CoV-1 stability also gave a neutral SNAP prediction. The statistical analysis showed that the correlation of SNAP scores in the five groups were significant with a P-value of <2e-16.

### Computational analysis of S viral mutations in SARS-CoV-1 isolates

This study analyzed mutations, found on the SARS-CoV-1 protein, which have been verified through experiments to have cellular or molecular effects on its functions. Analysis of SARS-CoV-1 isolates from the 2002-2003 and 2003-2004 outbreaks revealed the main driving mutations (32). Six residue positions of S protein, 344, 360, 472, 479, 480, and 487, were highlighted to have varying residues between the two isolates (32). The 2002-2003 isolate had residues K344, F360, L472, N479, D480, and T487. Meanwhile, the 2003-2004 isolate had the following corresponding residues R344, S360, P472, K479, G480, and S487. The effects of the residue changes from the 2002-2003 isolate to the 2003-2004 isolate are shown in Table 2. The residue change, N479K, would increase the binding affinity (ΔΔΔG = −1.008 kcal/mol) and reduced S stability (ΔΔG = 1.52 kcal/mol). Two mutations, L472P and T487S, decreased the binding affinity of S RBD-ACE2 by introducing ΔΔΔG at 1.875 kcal/mol and 0.752 kcal/mol, respectively. Furthermore, the mutation L472P destabilizes SARS-CoV-1 (ΔΔG = 1 kcal/mol). Therefore, understanding how the changes in these residues affect the stability and infectivity of the SARS-CoV-1 in the two isolates would help in the target of specific residues on SARS-CoV-2. A recent research to study the mutation of residues from the 2002-2003 isolates to residues in 2003-2004 isolate is consistent with this computational study (32).

**Table 2:** Computational prediction of the effect of residue change from 2002-2003 isolate to 2003-2004 isolate. (**N**)= Neutral, (**D**) = Decrease, (**I**) = Increase.

| SARS-CoV-1 |  | Energy Change |  |  | PolyPhen2 |  | SNAP |  |
| --- | --- | --- | --- | --- | --- | --- | --- | --- |
| 2002-2003 isolate | 2003-2004 isolate | Mutation | $\Delta\Delta\Delta G$(kcal/mol) | $\Delta\Delta G$(kcal/mol) | Score | Prediction | Score | Effect |
| K344 | R344 | K344R | 0 (N) | -0.02 (N) | 0 | Benign | -89 | Neutral |
| F360 | S360 | F360S | 0 (N) | 1.5 (D) | 0 | Benign | -77 | Neutral |
| L472 | P472 | L472P | 1.875 (D) | 1 (D) | 0.432 | Benign | 54 | Pathogenic |
| N479 | N479 | N479K | -1.008 (I) | 1.52 (D) | 0.598 | Possibly damaging | -80 | Neutral |
| D480 | G480 | D480G | -0.261 (I) | -0.25 (N) | 0.001 | Benign | -27 | Neutral |
| T487 | S487 | T487S | 0.752 (D) | -0.35 (N) | 0.003 | Benign | -89 | Neutral |

## Discussion

Coronaviruses have been the cause of the most recent pandemics. The most recent coronaviruses are SARS-CoV-1 and SARS-Cov-2. A recent study showed a close relationship between the sequence and the structure of SARS-CoV-1 and SARS-CoV-2 (33). The structural alignment performed in our study also revealed an evolutionary relationship between their S proteins, RBDs, and RBMs. However, the orientation of SARS-CoV-1 residues, S461 – N473, did not align properly with SARS-CoV-2 residues, A475 – N487. This imperfect alignment could be responsible for their varying binding affinities to the human hACE2 receptor (34).

The stability of the S protein is crucial for the rapid transmissions of infection (35). Understanding the role of mutations on S protein stability would help in designing therapeutic drugs and vaccines. Our prediction showed that more than half of the SARS-CoV-1 S mutations (~68%) destabilize the full-length S protein. Most of these destabilizing mutations involved the substitution of Glycine and Alanine residues, which are amino acids with hydrophobic side chains, with residues with longer hydrophobic side chains. Glycine has a short side chain which may hinder interactions with neighboring residues. Alanine has a deleted side chain which also makes it difficult to interact with neighboring residues. However, Alanine and Glycine exhibit hydrophobic effects which help to stabilize protein structures. The two most destabilizing missense mutations are A430W and A430F. The long side chains of Phenylalanine and Tryptophan disrupt the hydrophobic core of the SARS-CoV-1 S protein structure by introducing steric clashes. While most highly stabilizing mutations involved amino acids with polar side chains, such as Threonine, Arginine, and Serine, except for Glycine. The two most stabilizing missense mutations are G981W and T1059F (Fig 3B). Their polar side chains allow hydrogen bonding with water and ionic bond with nearby polar molecules. The prediction of the effects of mutations on the stability of SARS-CoV-1 S RBD revealed similar results with the analysis on full-length S protein. Among the top 5 positions with the highest average destabilizing effect, G418 and A430 were common (Fig 4A). Our study reveals a high correlation in the effects of mutations on the S RBD and the full-length S protein. This correlation shows that a mutation introduced into the SARS-CoV-1 will have a similar impact regardless of the parts, RBD or full-length S protein, used by the SARS-CoV-1. Due to the high similarity between SARS-CoV-1 and SARS-CoV-2, we compared the effects of mutations on their S proteins stabilities. After extrapolating the top results, we found out that the effects of mutations on protein stability are similar in corresponding positions on SARS-CoV-1 and SARS-CoV-2. However, D600G destabilizes SARS-CoV-1 S protein while D614G stabilizes SARS-CoV-2 S protein. We were also interested in looking at the residues that are different in corresponding positions. For instance, A430 on SARS-CoV-1 corresponds to S443 on SARS-CoV-2. The mutation A430S destabilizes SARS-CoV-1 S protein, while the mutation S443A stabilizes SARS-CoV-2 S protein.

The binding of the S protein to the ACE2 allows SARS-CoV-1 to enter the host’s cells(36). In this study, we predicted the effects of S mutations on the binding affinity between SARS-CoV-1 S RBD and hACE2. The top missense mutations decreasing binding affinity are T487Y, T487W, and G488P, with binding-energy changes (ΔΔΔG) of 20.018 kcal/mol, 13.42 kcal/mol, and 15.921 kcal/mol, respectively. These mutations occur on neighboring residues in the RBM of the SARS-CoV-1 S protein. By contrast, S432Y and N479M have the minimum binding-energy changes (ΔΔΔG) of −1.686 kcal/mol and −1.42 kcal/mol, respectively. Both mutations can strengthen the binding of SARS-CoV-1 S to hACE2. Furthermore, we looked at the Van der Waals interaction distance between interacting residues on SARS-CoV-1 S RBD and hACE2. The distance between residue T487 and K353 decreased when Threonine mutates to Tyrosine. On residue G488, all possible mutations will weaken its binding affinity for residues K353 and G354 on the hACE2 receptor. However, most missense mutations on SARS-CoV-1 N479 will strengthen its binding for H34 on hACE2 (Fig 6B – D). Compared to SARS-CoV-2 S RBD, the effects of mutations on the binding affinity of SARS-CoV-1 S RBD are similar. A few exceptions are in regions where the corresponding residues differ. For example, missense mutation S432V on SARS-CoV-1 decreased the binding energy by −0.22 kcal/mol. However, S432V has a neutral effect on its stability. Meanwhile, on SARS-CoV-2 S RBD, the missense mutation V445S does not affect its binding affinity.

SARS-CoV-1 modifies its S protein through N-linked Glycosylation, O-linked glycosylation, and palmitoylation. The S protein of SARS-CoV-1 possesses glycosylation sites similar to other coronaviruses(33). These modifications allow SARS-CoV-1 to bind differentially to hACE2 receptor, and to evade the immune system. Analysis of the effect of mutations on 20 N-Glycosylation sites and one palmitoylation site shows that these sites are crucial for the function of SARS-CoV-1. Furthermore, we used Polyphen2 and SNAP scores to predict the pathogenicity effects of the mutations, respectively. These tools have proven to identify non-synonymous substitutions with a high accuracy. In a study, Polyphen-2 achieved a prediction rates of 92% (37). In a different study, SNAP identified all neutral and non-neutral substitutions with an 80% accuracy (31). All five categories of the effects of mutations were significantly different, with P-value<2e-16. The Polyphen2 and SNAP scores of mutations with neutral effect were lower compared to the other four categories. The outcome of PolyPhen2 and SNAP predictions indicate the reliability of folding energy change (ΔΔG) in predicting the effect of missense mutations on the stability of SARS-CoV-1 S protein.

With our computational result, we compared viral isolates from 2002-2003 and 2003-2004 outbreaks. A previous study suggested that changes in residues affect the affinity of SAR-CoV-1 for hACE2 (32). In comparison, the order of affinity to hACE2 is 2002-2003 SARS-CoV-1 isolate > SARS-CoV-2 > 2003-2004 SARS-CoV-1 isolate (36). Therefore, we were curious to know the residue change within the RBD that resulted in a less severe 2003-2004 viral isolate. Our computational mutagenesis on the six residues in a previous study highlighted their role in hACE2 binding. The residues P472 and S487 weaken the binding affinity of the 2003-2004 viral isolate. The substitution of L472 with P472 in the 2003-2004 viral isolate reduces the binding of SARS-CoV-1 to L79 and M82 residues on hACE2. Threonine at position 487 has a stronger affinity to residues Y41, K353, and D355 on hACE2 than Serine at position 487. This is due to the methyl group in Threonine (32). Also, the residues S360, P472, and K479 destabilize the 2003-2004 viral isolate. Collectively, the effects of these changes in residues contribute to a less severe 2003-2004 viral isolate. However, the biological validation for effects of these viral mutations on protein stabilities and virus-receptor interactions are required. Other structural proteins such as, nucleocapsid (N), membrane (M), and envelope (E) protein play crucial roles in the function of coronaviruses. This computational saturated mutagenesis can be used to analyze these proteins for understanding SARS-CoV-1 and SARS-CoV-2. Particularly, our bioinformatic method provides a fast methodology to investigate all possible mutations and predict the potential dominant variants of coronaviruses in the future pandemics.

## Methods

### Structural preparation SARS-CoV-1 and SARS-CoV-2

We obtained the 3-dimensional structures of full-length S and RBD-hACE2 of both SARS-CoV-1 and SARS-CoV-2 from the RCSB Protein Data Bank(PDB) (38). The structure of a trypsin-cleaved SARS-CoV-1’s spike glycoprotein (PDB ID: 6ACG) was used for stability analysis. The structure of the complex of SARS-CoV-1 S RBD and the human hACE2 receptor (PDB ID: 2AJF) was used for stability and interaction analysis. For SARS-CoV-2, we obtained the protein complex structure of RBD-hACE2 (PDB ID: 6LZG) for stability and interaction analysis. The SARS-CoV-2 S (PDB ID: 6VYB) was used only for stability analysis. For structural alignment of RBD of SARS-CoV-1 and SARS-CoV-2, we used the structures PDB ID:2AJF and PDB ID:6M17, respectively. For structural alignment of the S protein of SARS-CoV-1 and SARS-CoV-2, we used the structures PDB ID:6ACG and PDB ID:6VYB, respectively. PyMOL (39) was used for the visualization of the PDB structures and for structural alignments.

### Computational mutagenesis and energy calculations

FoldX (20) was used for mutational analysis. We used the command line interface of FoldX to mutate each residue to the other 19 residues. The initial step used the ‘RepairPdb’ command to repair the wildtype protein structure. This was followed by either the use of the ‘BuildModel’ command and the ‘AnalyseComplex’ command to calculate the folding energy change and the interaction or binding energy change, respectively. For each mutation, we used FoldX to calculate the folding energy change (ΔΔG) and binding energy change (ΔΔΔG) (40).

The mathematical equation for the calculation of folding energy change (ΔΔG) is:

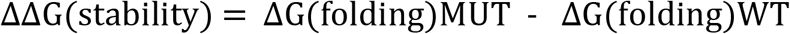

Theoretically, a negative ΔΔG means that the mutation leads to a more stable protein structure and a positive ΔΔG means that the mutation leads to a less stable protein structure. The five categories of the impact of the folding energy change (ΔΔG) are -highly stabilizing (ΔΔG<−2.0 kcal/mol), moderately stabilizing (−2.0<ΔΔG< −0.5 kcal/mol), neutral (0.5<ΔΔG<+0.5 kcal/mol), moderately destabilizing (+0.5<ΔΔG<2.0 kcal/mol), and highly destabilizing (ΔΔG > 2.0 kcal/mol).

The mathematical equation for the calculation of binding energy changes (ΔΔΔG) is:

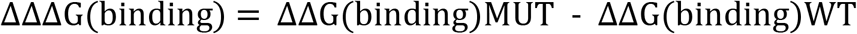

A negative ΔΔΔG means the mutation strengthens the binding energy and a positive ΔΔΔG means the mutation weakens the binding energy. The effect of the binding energy changes was also classified into five categories: large affinity decrease (ΔΔΔG> 0.5 kcal/mol), moderate affinity decrease (0.1<ΔΔΔG≤0.5), neutral (−0.1< ΔΔΔG≤0.1 kcal/mol), moderate affinity increase (−0.5< ΔΔΔG≤−0.5 kcal/mol), and large affinity increase (ΔΔΔG<−0.5 kcal/mol).

### Mutation pathogenicity and sequence-based Analysis

We used the Polymorphism Phenotyping v2 (PolyPhen2) (41) and Screening for non-acceptable polymorphisms (SNAP) (31) prediction tools to predict the pathogenicity of each missense mutations. We utilized the R programming language (https://www.r-project.org/.) for data visualization for the purpose of drawing inferences. Specifically, we constructed boxplots to compare the prediction of pathogenicity between PolyPhen2 and SNAP.

### Sequence and Structural similarity between SARS-CoV-1 and SARS-CoV-2

The FASTA sequences of SARS-CoV-1 and SARS-CoV-2 S proteins were retrieved from the Universal protein Knowledgebase (UniProtKB) (42). We performed the pairwise sequence alignment of SARS-CoV-1 (Entry: P59594) and SARS-CoV-2 (Entry: P0DTC2) using the Clustal Omega computer program (https://www.ebi.ac.uk/Tools/msa/clustalo/) and Jalview2 (www.jalview.org). We performed structural alignments of the 3-D structures of SARS-CoV-1 and SARS-CoV-2 using PyMOL (http://www.pymol.org/). The “fetch” and the “align” commands on PyMOL aligned the single chains of the spike proteins of SARS-CoV-1 (PDB ID: 6ACG, chain A) and SARS-CoV-2 (PDB ID: 6VYB, chain A), and RBD of SARS-CoV-1 (PDB ID: 2AJF, chain E) and SARS-CoV-2 (PDB ID: 6M17, chain E).

## Conclusions

Saturated computational mutagenesis of SARS-CoV-1 S protein proved to be effective in analyzing energy changes. Missense mutations in key residues such as A430 and S500 stabilized and destabilized SARS-CoV-1 full-length S and RBD, respectively. Moreover, missense mutations on residues G488 and T487 weakened the binding affinity of SARS-CoV1 S to hACE2. Mutation pathogenicity analysis showed that most highly destabilizing and highly stabilizing missense mutations would have a damaging effect on the SARS-CoV-1 S function. We also showed that missense mutations on N-linked glycosylation sites would stabilize SARS-CoV-1 S. The analysis of viral isolates from 2002-2003 and 2003-2004 showed that residue changes N479K, L472P, and F360S destabilized the S protein of 2003-2004 viral isolate leading to a reduction in infection rate. Also, most of the S missense mutations on SARS-CoV-1 had a similar stabilizing or destabilizing effect on corresponding residues on SARS-CoV-2. This approach can provide large-scale mutagenesis for future experimental studies on the coronavirus research.

## Acknowledgements

We thank the Howard University Junior Faculty Writing & Creative Works Summer Academy, Ernest E. Just-Percy L. Julian Graduate Research Assistantship and Research Centers for Minority Institutions (RCMI).

## Author contributions

ST designed the study and supervised the study. AS and ST performed the bioinformatics analysis. AS, VM, RR, DW, DL, LL, QT and ST analyzed data. AS and ST wrote the manuscript, and all authors critically revised the manuscript.

